# p38-MAPK is prerequisite for the synthesis of SARS-CoV-2 protein

**DOI:** 10.1101/2023.09.27.559660

**Authors:** Priyasi Mittal, Nitin Khandelwal, Yogesh Chander, Assim Verma, Ram Kumar, Chayanika Putatunda, Sanjay Barua, Baldev Raj Gulati, Naveen Kumar

## Abstract

The inhibition of p38 mitogen-activated protein kinase (p38-MAPK) by small molecule chemical inhibitors was previously shown to impair severe acute respiratory syndrome coronavirus 2 (SARS-CoV-2) replication, however, mechanisms underlying antiviral activity remains unexplored. In this study, reduced growth of SARS-CoV-2 in p38-α knockout Vero cells, together with enhanced viral yield in cells transfected with construct expressing p38α, suggested that p38-MAPK is essential for the propagation of SARS-CoV-2. The SARS-CoV-2 was also shown to induce phosphorylation (activation) of p38, at time when transcription/translational activities are considered to be at the peak levels. Further, we demonstrated that p38 supports viral RNA/protein synthesis without affecting viral attachment, entry, and budding in the target cells. In addition, we demonstrated that long-term culture of SARS-CoV-2 in the presence of p38 inhibitor SB203580 does not easily select resistant viral mutants. In conclusion, we provide mechanistic insights on the regulation of SARS-CoV-2 replication by p38 MAPK.

## Introduction

Severe acute respiratory syndrome coronavirus 2 (SARS-CoV-2) which emerged in 2019 has resulted in more than 767.5 million infections and 6,947,192 fatalities as of September 2023. (WHO 2023). Vaccine is available but emergence of the antigenic variants has raised concerns about the immunity induced due to vaccination or prior infection (Zhao et al., 2004).

Remdesivir, Favipiravir, Itolizumab, Tocilizumab, Hydroxychloroquine, and Ivermectin are among antiviral medications used to treat COVID-19. These medications decrease the number of patients hospitalized and the disease severity of the condition (Chen et al., 2020; Food and Administration, 2020; Team et al., 2020; Warren et al., 2016). However specific and effective antiviral medication against COVID-19 are still lacking.

Till date, more than 184 antiviral drugs have been approved by the Food and Drug Administration (FDA) to combat with viral infections. The mechanism of action of most of these approved drugs is based on directly targeting viral protein expression (Chaudhuri et al., 2018). However, even when successful, the drugs can eventually fail because of the emergence of drug-resistant mutants (Hovi et al., 2002; Irwin et al., 2016; Li and Chung, 2019; Locarnini and Bowden, 2010; Pillay and Zambon, 1998).

To facilitate replication, viruses interact with numerous factors by RNA-protein, protein-protein and protein-lipid interactions (Merino-Ramos et al., 2016). Upon infection, viruses exploit the host cells by subverting the host factors, remodeling subcellular membranes, modulation of cellular proteins and ribonucleoprotein complexes and usurping cellular metabolic pathway (Watanabe et al., 2001). Recent progress through transcriptomics studies, genome-wide knockdown/knockout studies have enabled identification of numerous cellular factors which are essential for virus replication (Venter et al., 2001). The host cell factors which are critically required for virus replication but are dispensable for the host may be targeted for antiviral drug development (Chander et al., 2021b; Kumar et al., 2018a; Kumar et al., 2018c). Since the genetic variability of the host is quite low as compared to the viral genome, host-directed therapies should have fewer tendencies in inducing antiviral drug resistance.

Recently we demonstrated that p38 MAPK inhibitor SB203580 block SARS-CoV-2 yield in Vero cells (In-press, paper entitled p38 MAPK Inhibitor SB203580Suppresses SARS-CoV-2 Replication in Annals of biology, Ref. No. AOB/2023/14). However, the precise mechanism of the antiviral action of action remains elusive. In this study we provided mechanistic insights on the regulation of SARS-CoV-2 replication by p38-MAPK, along with studying development of drug-resistant SARS-CoV-2 mutants under long-term selection pressure of p38 inhibitor.

## Materials and Methods

### Cells and viruses

Vero (African green monkey kidney cells), T-293 (human embryonic kidney) and p38- α MAPK knockout Vero cells available at National Centres for Veterinary Type Culture (NCVTC), Hisar were grown in Dulbecco’s Modified Eagle’s Medium high glucose (Lonza, cat number-12-604F BE12- 604F, USA) supplemented with antibiotics and 10% heat-inactivated fetal bovine serum (FBS) (D6429, Sigma, St. Louis, USA) and antibiotic (Penicillin-Streptomycin-Amphotericin B Suspension 100X) (A5955, Sigma, USA). Wild type (SARS-CoV-2/Human-tc/India/2020/Hisar-4907) SARS-CoV-2 (VTCC Accession Number of VTCCAVA 294 (SARS-CoV-2/India/2020/tc/Hisar/4907) were available at NCVTC, Hisar. Virus was propagated in Vero cells in the Biosafety level 3 (BSL-3) laboratory of ICAR-National Research Centre on Equines (NRCE), Hisar, India (Khandelwal et al., 2021). The virus was quantified by plaque assay and viral titres were measured as plaque forming unit per millilitre (PFU/ml) (Kumar et al., 2021).

### Inhibitor

Adezmapimod (SB203580), is a pyridinyl imidazole inhibitor were procured from Biogems (1524762, Ariano Irpino, Italy). These inhibitors were dissolved in Dimethyl sulfoxide (DMSO), thereby DMSO was used as a vehicle control in the experiments (Henklova et al., 2008)

### Antibodies

SARS-CoV-2 Nucleocapsid antibody (produced in mouse) as a primary antibody was procured from (MA5-35943, Invitrogen, 1:1000-1:10,000), Rabbit phospho-p38 MAPK and total p38 MAPK Antibody were procured from (9211S and 9212S, Cell Signalling technology, Massachusetts, USA). Mouse anti-β actin primary antibody, Goat Anti-mouse IgG HRP conjugate (665739 /Merck/Massachusetts, USA/1: 10000) Goat Anti-Rabbit IgG HRP conjugate (632131/Merck/Massachusetts, USA, 1:10,000 were procured from Merck (USA). Mouse ß-Actin (3700S/ 1:1000/Cell Signalling technology/Massachusetts, USA) used as primary antibody

### Overexpression of p38-α to rescue the inhibitory effect of p38-α depletion SARS-CoV-2

Plasmid constructs expressing p38 MAPK (pDEST40-p38-α) was available at NCVTC, Hisar. To further confirm the SARS-CoV-2 supportive role of p38-α, T-293 cells were either transfected with pDEST40-p38-α (5 µg) or with the empty vector (control) using Lipofectamine 3000 (Thermo Fisher Scientific, Waltham, MA, USA). At 24 h post-transfection, the cells were infected with SARS-CoV-2 at MOI of 10 and the virus released in the infected cell culture supernatant at 24 hpi was quantified by Quantitative Real time PCR (qRT-PCR).

### Attachment Assay

Confluent monolayer of Vero cells, in triplicates, were pre-incubated with SB203580 (2.5µg/ml) or vehicle control for 30 minutes followed by SARS-CoV-2 infection (MOI of 5) at 4□C for 1.5h. Thereafter, the cells were washed with cold ice PBS and lysates were prepared by freeze-thaw. Virus attached to cell in presence and absence of inhibitor is quantified by plaque assay (Kumar et al., 2022).

### Entry assay

The confluent monolayer of Vero cells, in triplicates were prechilled at 4 °C and infected with SARS-CoV-2 (MOI-5) at 4□C for 1 h which allowed the virus attachment to the cells and restricted the viral entry. Thereafter, the cells were washed with ice-cold PBS and add inhibitor (SB203580, 2.5µg/ml) or vehicle controls were added, and cells were incubated at 37^δ^C for 1 h to permit the viral entry. Cells were again washed with PBS, followed by addition of DMEM without any inhibitor. The cells were further incubated at 37□C. Supernatant was collected at 16 hpi and the infectious virus particles were quantified by plaque assay (Kumar et al., 2019a).

### Virus Release Assay

Confluent monolayer of Vero cells, in triplicates, were infected with MOI of 5 for 1 h at 37□C followed washing with PBS and addition of fresh DMEM. At 8 hpi cells when virus presumably starts budding, cells were again be washed with PBS followed by addition of fresh medium having inhibitor (2.5 µg/ml) or vehicle control. Supernatant is collected at 0.5 h and 1-hour post-drug treatment and the virus released is quantified by plaque assay.

### qRT-PCR

Confluent monolayer of Vero cells was infected with SARS-CoV-2 (MOI=5) for 1 h at 37□C followed by washing with cold ice PBS and addition of fresh DMEM. At 3 hpi, when early stages of life cycles like attachment, entry and uncoating had occurred and RNA synthes is likely to start, 2.5 µg/ml of SB203580 or 0.05% DMSO was added. Cells were scraped at 18 hpi to quantify the viral gene and house-keeping control gene (GAPDH) by qRT-PCR. All the values were normalized with GAPDH housekeeping control gene. Relative fold-change in viral RNA copy number were determined by ΔΔ Ct method (Kumar et al., 2016).

### Effect on the synthesis of viral proteins

Confluent monolayer of Vero cells in 30 cm tissue culture dishes were infected with SARS-CoV-2 at MOI of 5 followed by addition of SB203580 (2.5µg/ml) (at 3 hpi). The cells were scrapped at 24 hpi and cell lysate were prepared in RIPA buffer. The levels of viral and house-keeping control proteins were analyzed in Western blot. SARS-CoV-2 Nucleocapsid antibody (produced in mouse) was procured from (MA5-35943/Invitrogen/Massachusetts, USA/1:1000-1:10,000) used as primary antibody. Goat Anti-mouse IgG HRP conjugate (665739 /Merck/Massachusetts, USA/1: 10000) and its substrate (DAB) were used as manufacturer’s protocol.

### Activation of p38 MAPK (p38 Phosphorylation

Confluent monolayers of Vero cells in 100 cm tissue culture dishes were infected with SARS-CoV-2 at MOI of 10 for 1 h, followed by washing with ice cold PBS and addition of fresh DMEM. Cells were scrapped off at 1hpi, 4hpi, 8hpi and 12hpi cell lysates were prepared. The levels of phosphorylated p38 MAPK, total p38 MAPK and house-keeping (beta-actin) control proteins were analyzed in Western blot.

### Selection of SB203580-resistant virus variants

Vero cells were infected with SARS-CoV-2 at MOI of 0.1 in the medium containing 0.05% DMSO or 0.5 µg/mL of SB203580. At 36-48 hpi, supernatant was collected from the virus infected cells (named passage 1 [P1]) and quantified by plaque assay. Fifty such sequential passages were carried out. The original virus stock (P0), P50-SB203580 and P50-control viruses were again used to infect Vero cells at an MOI of 0.1 with either 2.5 µg/mL SB203580 or 0.05% DMSO. At 36-48 hpi, viral titers in the infected cell culture supernatant were quantified by plaque assay.

## Results

### p38 MAPK is essential for SARS-CoV-2 replication

To evaluate the role of p38 MAPK on SARS-CoV-2 replication, we measured yield of SARS-CoV-2 in WT- and p38-knockout (KO) Vero cells. As shown in **Fig. 1A**, p38-knockout cells had significantly lower titer as compared to the Wt Vero cells, suggesting p38 MAPK supports SARS-CoV-2 replication.

**Fig 1:**
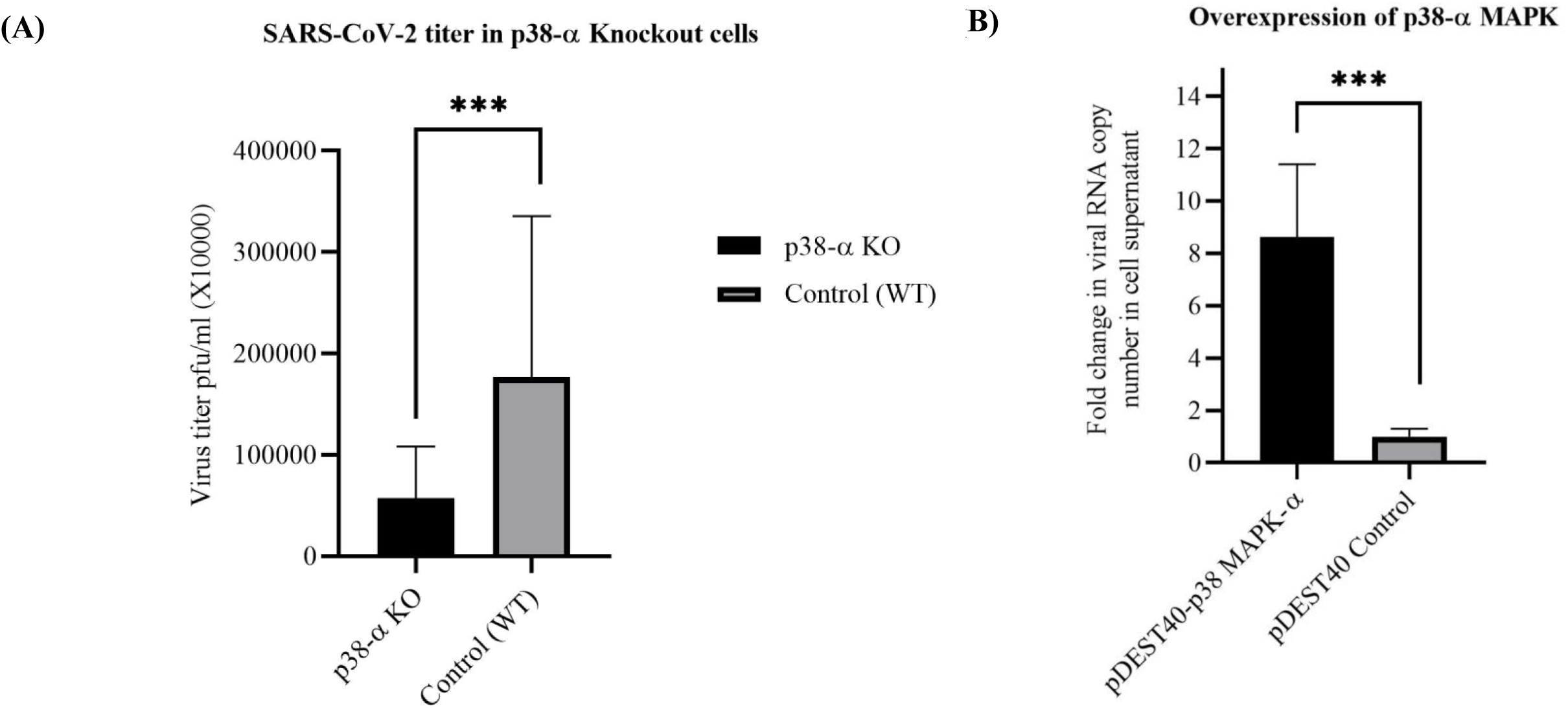
p38 is prerequisite for the propagation of SARS-CoV-2. ***(A) Growth of SARS-CoV-2 in p38-#x03B1; KO cells*.** The confluent monolayers of WT Vero cells and p38 knockout Vero cells, in triplicates, were infected with SARS-CoV-2 at MOI 1. At 24 hpi, supernatant was collected and yields of infectious progeny virus particles were determined by plaque assay. Values are means ± SD and representative of the result of at least 3 independent experiments. Pair-wise statistical comparisons were performed using Student’s t test (***= P <0.001). ***(B) Overexpression of p38.*** 293T cells were transfected with plasmid construct that express p38 MAPK or with the empty vector. At 24 h post-transfection, the cells were be infected with SARS-CoV-2 at MOI of 10 and the virus released in the infected cell culture supernatant at 24 hpi was quantified by qRT-PCR. Threshold cycle (Ct) values was analysed to determine relative fold-change in copy numbers of total RNA and mRNA. Values are means ± SD and representative of the result of at least 3 independent experiments. Pair-wise statistical comparisons were performed using Student’s t test (***= P <0.001).

To further confirm, we measured the yield of SARS-CoV-2 in 293T cells transfected with the construct that express p38 (pDEST40-p38-α). As shown in **Fig 1B**, the cells transfected with p38 expressing protein had significantly higher titre as compared to the cells that received empty vector, which further confirmed that p38 is essential for the propagation of SARS-CoV-2.

### p38 MAPK inhibition impairs SARS-CoV-2 RNA and protein synthesis

We performed virus step-specific assays to evaluate the role of p38 MAPK in SARS-CoV-2 life cycle. The p38 inhibitor did not affect SARS-CoV-2 attachment (Fig 2a), entry (Fig 2b) and budding (Fig 2c) in the target cells.

**Fig 2:**
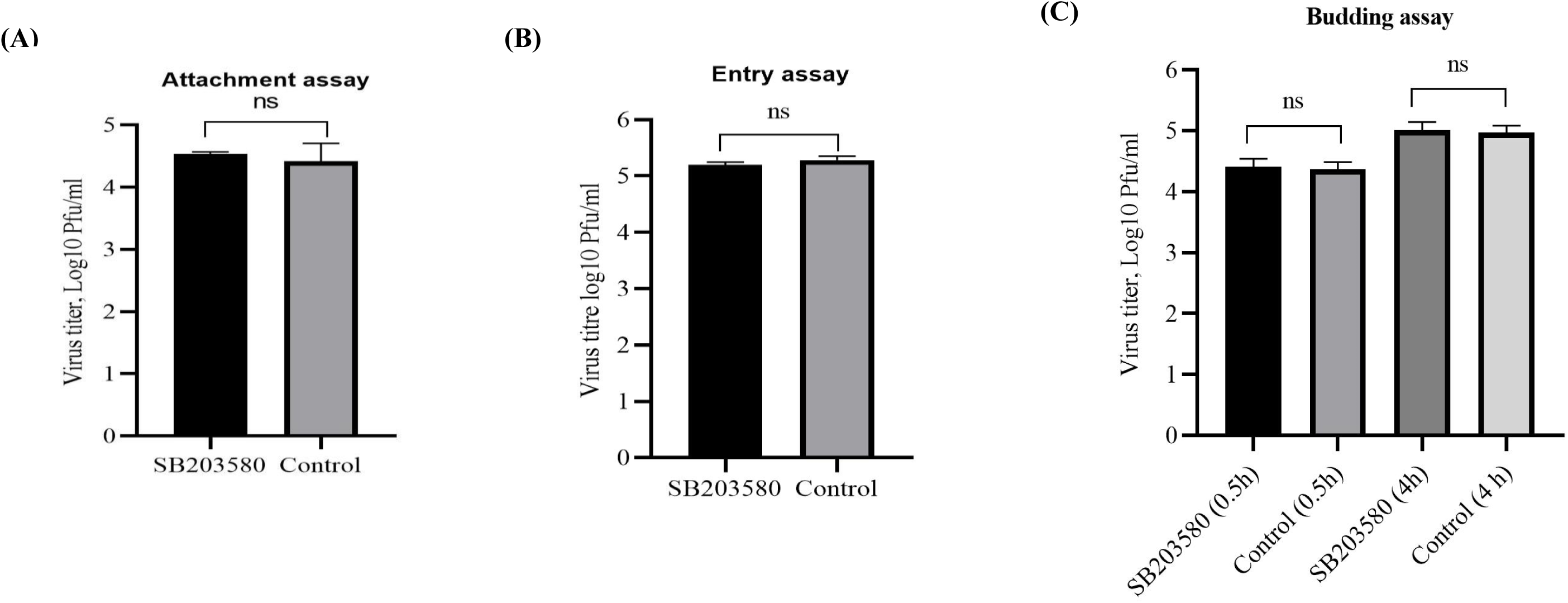
p38 inhibitor does not inhibit virus attachment, entry and budding. ***(A) Attachment assay*.** Vero cells were pre-incubated with SB2203580 or vehicle control for 30 minutes followed by SARS-CoV-2 infection (MOI 5) at 4□C for 1.5 h. The cells were then washed 5 times with PBS and the cell lysates were prepared by rapid freeze-thaw method. Virus attached to the cells in the presence of inhibitor or vehicle control was quantified by Plaque assay (ns= non-significant). *(**B) Entry assay**.* Confluent monolayer of Vero cells, in triplicates were infected with SARS-CoV-2 infection (MOI 5) at 4□C for 1 h, followed by washing with ice-cold PBS. The attached virus was allowed to enter at 37□C in presence of inhibitor or vehicle control. The virus released in the supernatant was quantified at 16 hpi by plaque assay. ***(C) Budding assay***. Confluent monolayer of Vero cells was infected with SARS-CoV- 2 at MOI 5 for 1 h followed by washing with PBS. At 8 hpi, cells were again washed with PBS and fresh medium with inhibitor was added. Supernatant was quantified by plaque assay.

To determine the effect of p38 inhibition on viral RNA/protein synthesis, SB203580 was applied at 3 hpi, a time when early steps of SARS-CoV-2 life cycle (attachment, entry) are expected to occur. As shown in **Fig 3A**, as compared to the Control, there was significantly less RNA in cells treated with SB203580, which suggested that p38 may be required for efficient synthesis of SARS-CoV-2 RNA. Like RNA, we also observed significantly less SARS-CoV-2 proteins in cells treated with SB203580, as compared to the control-treated cells **(Fig 3B, 3C)**.

**Fig 3:**
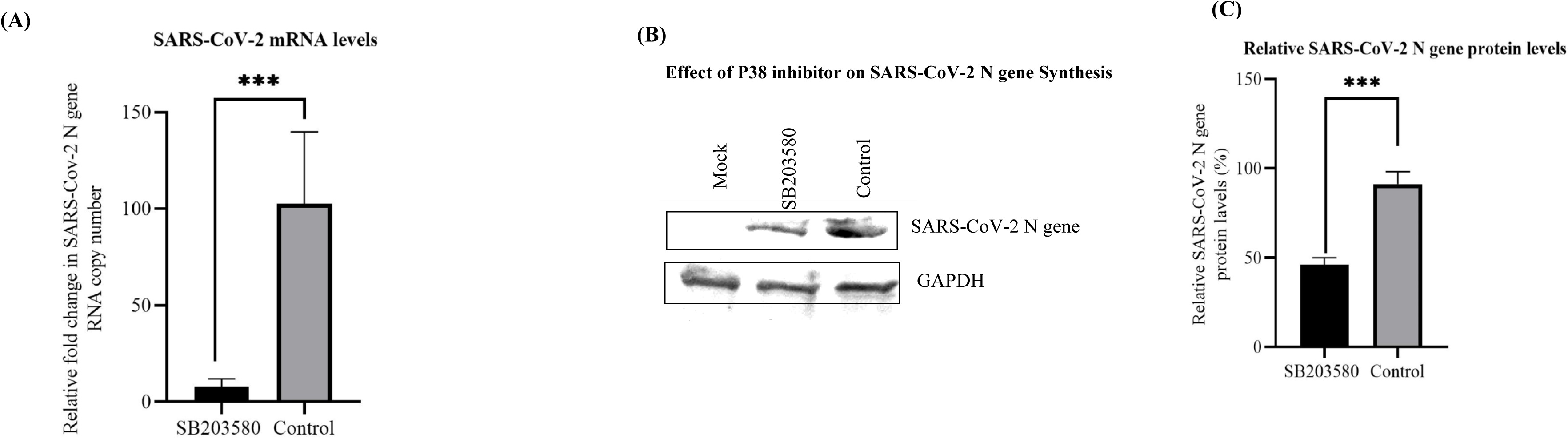
Effect of SB203580 on levels of viral RNA and protein. ***(A) RNA level*.** Confluent monolayers of Vero cells, in triplicates, were infected with SARSCoV-2 for 1 h at MOI 5. SB203580 was added at 3 hpi and cells were harvested at 12 hpi to determine the levels of SARS-CoV-2 RNA by qRT-PCR. Threshold cycle (Ct) values were analysed to determine relative fold-change in copy numbers of total RNA and mRNA. Values are means ± SD and representative of the result of at least 3 independent experiments. Pair-wise statistical comparisons were performed using Student’s t test (***= P <0.001). ***(B) Protein levels.*** Confluent monolayers of Vero cells were infected with SARS-CoV-2 at an MOI of 5. The inhibitor or DMSO was applied at 3 hpi and the cells were scrapped at 24 hpi to examine the levels of viral proteins by immunoblotting. ***(C) Quantitation of protein levels.*** The blots were quantified by densitometry (ImageJ) and the data are presented as mean with SD. The levels of viral proteins (upper panel), along with housekeeping GAPDH protein (lower panels) is shown. Pair-wise statistical comparisons were performed using Student’s t-test. ***=P<0.001. Values are means ±SD and representative of the result of at least tree-independent experiments.

### SARS-CoV-2 induces activation of p38 MAPK

To further examine whether the SARS-CoV-2 induces p38 activation (phosphorylation), the cell lysate was collected at different time points and subjected to western blotting analysis. There were minimal or no detectable level of p-p38 in mock-infected cells as well as in cell lysates collected at 1 hpi, 4 hpi, 8 hpi and 12 hpi. The highest level of p-p38 was observed at ∼ 8 hpi **(Fig. 4A, 4B)**, a time when transcription and translation is expected to occur. The total level of p38 as well as level of ß-actin (housekeeping gene) was at comparable level in all the samples **(Fig. 4A)**. This suggested that SARS-CoV-2 induces activation is coupled with transcription/translation of SARS-CoV2 transcripts.

**Fig 4:**
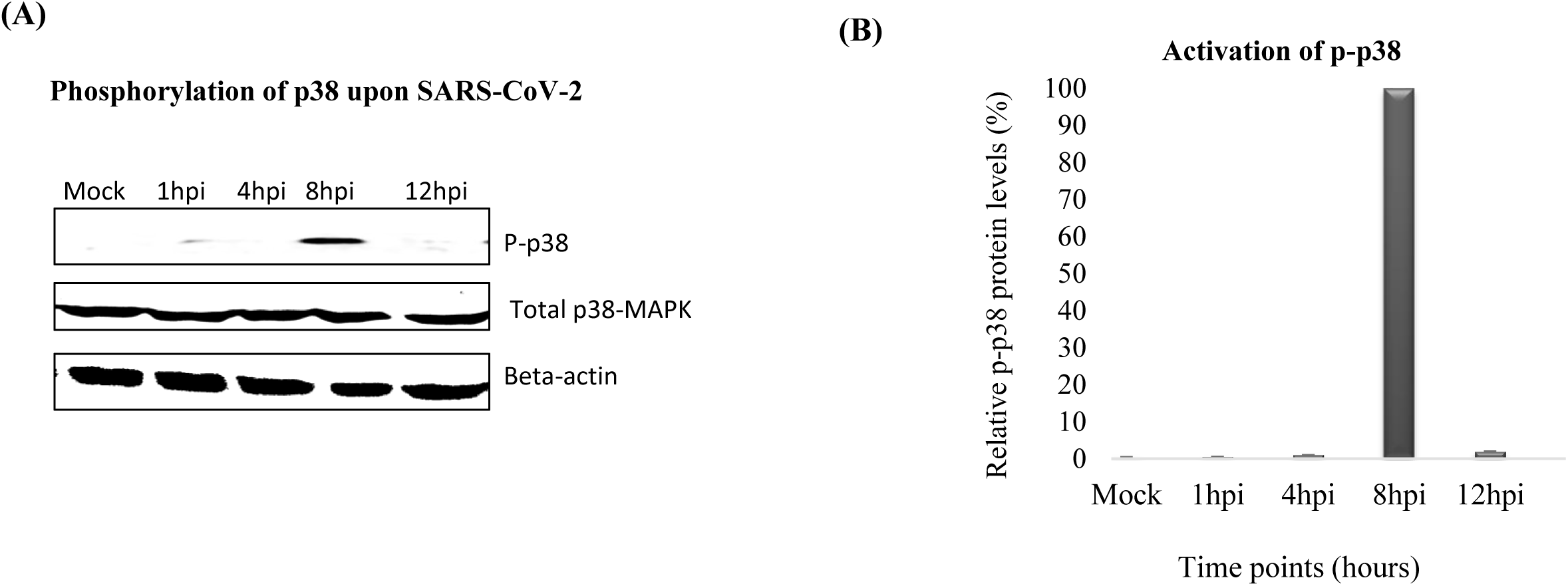
SARS-CoV-2 activates p38 MAPK. Confluent monolayers of Vero cells were infected with SARS-CoV-2 at MOI of 10 for 1 h, followed by washing with ice cold PBS and addition of fresh DMEM. Cell was scrapped off at 1 hpi, 4 hpi, 8 hpi and 12 hpi in PBS to prepare the cell lysates. The levels of p38 phosphorylation at different time points and house-keeping control proteins were analysed in Western blot (immunoblotting) **(A).** The levels of phosphorylation-p38 (upper panel), along with total p38 MAPK (middle panels) and housekeeping beta-actin gene (lower panels) are shown. The blots were quantified by densitometry (ImageJ) and the data are presented as mean with SD **(B).**

### Selection of potential Drug-Resistant virus variants

In order to evaluate the development of drug-resistant virus variants, SARS-CoV-2 was serially passaged 50 times in the presence of SB203580 or vehicle-control (DMSO). At P50, SARS-CoV-2-P50-SB203580 and SARS-CoV-2-P50-control, together with the original virus (SARS-CoV-2-P0) were evaluated for their sensitivity to SB203580. The magnitude of the suppression of virus yield in SB203580-treated cells was at lower level in SARS-CoV-2-P50-SB203580, as compared to SARS-CoV-2-P50-control, which suggested the selection of resistant mutants by SB203580 **(Fig. 5A)**. However, P0 virus replicated ∼46 fold higher as compared to (P60-SB203580) and nearly ∼42 fold as compared to (P60-Control) **(Fig 5B)**, which could be due to higher fitness due to high sequential passage. Nevertheless, a complete resistant could not be achieved.

**Fig 5:**
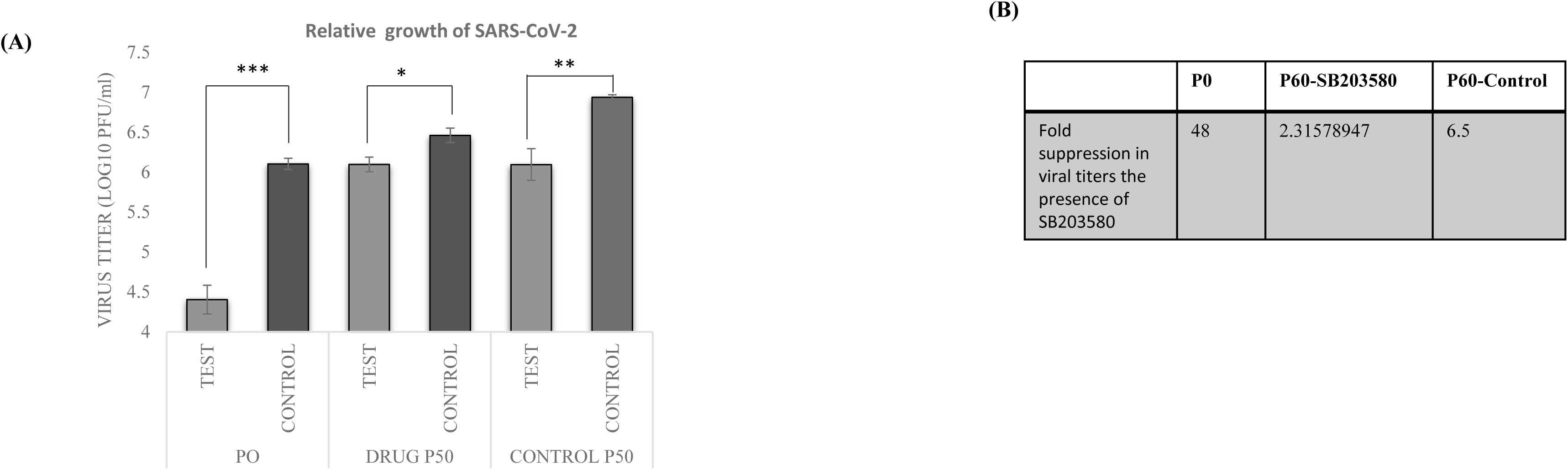
Selection of potential drug-resistant virus variants. Vero cells were infected with SARS-CoV-2 at MOI of 0.1 and grown in the presence of either 0.5 μg/ml of SB203580 or vehicle control (0.05% DMSO). The progeny virus particles released in the supernatant were harvested either at 36-48 hpi or when ∼75% cells exhibited CPE. Fifty (50) such sequential passages were made***. (A). Sensitivity of P50- SB203580-SARS-CoV-2 and P50-Control-SARS-CoV-2 to SB203580:*** Vero cells, in triplicate, were infected with P0, P50-SB203580, or P50-Control passaged viruses at MOI of 0.1 in the presence of either 2.5 μg/ml SB203580 or 0.05% DMSO, and the progeny virus particles released in the supernatant at 24- 36 hpi were quantified. Fold change were determined. Error bars indicate SD. Pair-wise statistical comparisons were performed using Student’s t-test. (***=P<0.001). ***(B)*** Fold-suppression of virus yield is also shown

## Discussion

Drug repositioning or repurposing is an effective strategy to immediately respond to the emerging diseases (Becker et al., 2021). Based on virtual screen and *in vitro* effects, several therapeutics drugs were repurposed to treat COVID19. These includes, Remdesivir, Favipiravir, Itolizumab, Tocilizumab, Hydroxychloroquinine and Ivermectin (Rajnik et al., 2021; ul Qamar et al., 2020). These medications decrease the number of patients hospitalized and the severity of the condition (Chen et al., 2020; Food and Administration, 2020; Team et al., 2020; Warren et al., 2016). However specific and effective antiviral medication against COVID-19 is lacking.

An alternative approach to developing antiviral strategies which has been minimally explored is to design drugs that target host cell proteins needed for virus replication. The family members of mitogen-activated protein kinase (MAPK) are the key kinases involved in most signal transduction pathways (Cargnello and Roux, 2011; Cohen, 1997; Kumar et al., 2019a). p38 is usually activated in response to stress therefore, it is also considered as a stress-activated MAPK (Koul et al., 2013). p38 has four splice variants (isoforms) which include p38α (MAPK14), p38β (MAPK11), p38γ (MAPK12) and p38δ (MAPK13) (Huang et al., 2009). A wide variety of viruses are known to directly interact with p38 or its substrates (Maik-Rachline et al., 2020). While some of the interactions are proviral others are inhibitory to virus replication. The previous studies have demonstrated that inhibitor targeting p38 MAPK suppresses SARS-CoV-2 replication (Higgins et al., 2021). However, the mechanisms underlying regulation of SARS-CoV-2 replication remains elusive. In this study, we provided the mechanistic insights on regulation of SARS-CoV-2 replication by p38 MAPK.

In order to examine the role of p38 signaling in SARS-CoV-2 infections, initially we propagated SARS-CoV-2 in p38 MAPK knockout cells. The reduction in the virus yield p38-depleted (CRISPR/Cas9 knockout) cells and enhanced growth in cells transfected with the plasmid construct that expresses p38-□ MAPK suggested that p38 MAPK signaling is a prerequisite for SARS-CoV-2 replication. Further, it was also demonstrated that SB203580 inhibitor-mediated suppression of SARS-CoV-2 replication is primarily due to the reduced levels of viral proteins and partly due to reduced levels of viral RNA but without any significant effect on viral attachment, entry, and budding. Our finding are in agreement with other studies on wherein p38 MAPK was shown to promote synthesis of viral proteins (Banerjee et al., 2002; Cook, 2016; Higgins et al., 2021; Mudaliar et al., 2021; Su et al., 2017; Sugasti-Salazar et al., 2021; Zhan et al., 2020). However p38 may also supports RNA synthesis [respiratory syncytial virus (RSV) and influenza A virus mRNA synthesis (Choi et al., 2016), viral assembly HCV (Cheng et al., 2020) which seems to be due to the involvement of different downstream effector molecules (Chander et al., 2021a; Cuadrado and Nebreda, 2010).

A major risk factor in death of COVID-19 patients is hyper-induction of cytokine secretion. Since the blockade of p38 also dampens virus induced cytokine storm (Börgeling et al., 2014; Chander et al., 2021a; Du et al., 2016; Galván Morales et al., 2014; Griego et al., 2000; Lee et al., 2007; Mikkelsen et al., 2009; Peng et al., 2014; Shapiro et al., 1998), the drugs targeting p38 may have dual effects, first in limiting virus production in the target cells and secondly, in dampening the cytokine storm.

Due to mutations at the drug-binding sites, viruses rapidly acquire drug-resistant variants, therefore, developing antiviral therapeutics is a major challenge (Chaudhary et al., 2015; Kumar et al., 2014; Kumar et al., 2011b; Pawlotsky, 2012). Whereas directly acting antiviral agents rapidly induce generation of drug-resistant viral mutants (Kumar et al., 2011b), host-directed therapies are less prone to generate drug-resistant mutants (Chaudhary et al., 2015; Kumar et al., 2019b; Kumar et al., 2011a; Kumar et al., 2020; Kumar et al., 2008; Kumar et al., 2018b; Kumar et al., 2018d; Xu et al., 2020) because viruses cannot easily regain the missing cellular functions by mutation (Kumar et al., 2020; Kumar et al., 2018d). However, recent evidence suggests that resistance to host-directed antiviral agents can occur at a relatively low magnitude upon long-term limiting availability of the targeted cellular factor (Hopcraft and Evans, 2015). To evaluate the development of SB203580-resistant mutants, SARS-CoV-2 was serially passaged 50 times in the presence or absence of SB203580. As compared to the P50-Contol, P50- SB203580 virus replicated at significantly higher titres in SB203580-treated cells, suggesting the emergence of SB203580-resistantSARS-CoV-2 mutants. However, a complete resistance could not be achieved. Previous study by our group on buffalopox virus suggests that virus switches to use alternate cellular factor (p38-α to p38-γ) upon long-term restricted availability of the inhibitor targeting p38-α (Chander et al., 2022).

Since host-directed agents interfere with the host cell metabolism, their use may be associated with side effects (Lamarche et al., 2012). However, the large number of the host-directed agents licensed to treat cardiovascular and inflammatory diseases or cancers have minimal or no adverse side effects (Shepherd et al., 2005; Zeisel et al., 2013). However, further validation and *in vivo* efficacy of SB203580 in COVID-19 patients is essential before actually introducing it from the research into the clinical settings. In conclusion, p38 MAPK serves as a essential cellular factor for the synthesis of SARS-CoV-2 proteins, and may serve as a novel target for antiviral drug development against COVID19. SARS-CoV-2 does not easily select SB203580-resisatnt viral mutants. However, under long-term selective pressure of p38 inhibitors, a partial resistance may be developed.

### Data availability statement

The original contributions presented in the study are included in the article further inquiries can be directed to the corresponding authors.

### Author contributions

Naveen Kumar contributed to the study conception and design. Material preparation was performed by Priyasi Mittal, Nitin Khandelwal, Yogesh Chander, Assim Verma, Ram Kumar, Garvit Kumar. Data analysis was performed by Priyasi Mittal, Baldev Raj Gulati, Naveen Kumar, Chayanika Putatunda and Sanjay Barua. The first draft of the manuscript was written by Priyasi Mittal and Naveen Kumar, and all authors commented on previous versions of the manuscript. All authors read and approved the final manuscript.

## Funding

This work was supported by Indian Council of Agricultural Research, New Delhi (grant number IXX14586 to N-Ku and NASF/ABA-8027/2020-21 to N-Ku and B.R.G.)

## Conflict of interest

The authors declare that the research was conducted in the absence of any commercial or financial relationships that could be construed as a potential conflict of interest

## Acknowledgments

This work was supported by the Science and Engineering Research Board, Department of Science and Technology, Government of India (grant number CVD/2020/000103, CRG/2018/004747 and CRG/2019/000829 to N. Kumar and S. Barua). A part of this study belongs to the PhD thesis work of Priyasi Mittal

## Notes

### Competing Interest Statement

The authors have declared no competing interest.

## References

Banerjee, S, Narayanan, K, Mizutani, T, and Makino, S (2002): Murine coronavirus replication-induced p38 mitogen-activated protein kinase activation promotes interleukin-6 production and virus replication in cultured cells. Journal of virology 76, 5937–5948.

Becker, M, Dulovic, A, Junker, D, Ruetalo, N, Kaiser, PD, Pinilla, YT, Heinzel, C, Haering, J, Traenkle, B, and Wagner, TR (2021): Immune response to SARS-CoV-2 variants of concern in vaccinated individuals. Nature communications 12, 3109.

Börgeling, Y, Schmolke, M, Viemann, D, Nordhoff, C, Roth, J, and Ludwig, S (2014): Inhibition of p38 mitogen-activated protein kinase impairs influenza virus-induced primary and secondary host gene responses and protects mice from lethal H5N1 infection. Journal of Biological Chemistry 289, 13–27.

Cargnello, M, and Roux, PP (2011): Activation and function of the MAPKs and their substrates, the MAPK-activated protein kinases. Microbiology and molecular biology reviews 75, 50–83.

Chander, Y, Kumar, R, Khandelwal, N, Singh, N, Shringi, BN, Barua, S, and Kumar, N (2021a): Role of p38 mitogen-activated protein kinase signalling in virus replication and potential for developing broad spectrum antiviral drugs. 31, 1–16.

Chander, Y, Kumar, R, Khandelwal, N, Singh, N, Shringi, BN, Barua, S, and Kumar, N (2021b): Role of p38 mitogen□activated protein kinase signalling in virus replication and potential for developing broad spectrum antiviral drugs. Reviews in Medical Virology 31, 1–16.

Chander, Y, Kumar, R, Verma, A, Khandelwal, N, Nagori, H, Singh, N, Sharma, S, Pal, Y, Puvar, A, and Pandit, R (2022): Resistance evolution against host-directed antiviral agents: Buffalopox virus switches to use p38-□ under long-term selective pressure of an inhibitor targeting p38- α. Molecular Biology and Evolution 39, msac177.

Chaudhary, K, Chaubey, KK, Singh, SV, and Kumar, N (2015): Receptor tyrosine kinase signaling regulates replication of the peste des petits ruminants virus. Acta Virol 59, 78–83.

Chaudhuri, S, Symons, JA, and Deval, J (2018): Innovation and trends in the development and approval of antiviral medicines: 1987–2017 and beyond. Antiviral research 155, 76–88.

Chen, H, Zhang, Z, Wang, L, Huang, Z, Gong, F, Li, X, Chen, Y, and Wu, JJ (2020): First clinical study using HCV protease inhibitor danoprevir to treat COVID-19 patients. Medicine 99.

Cheng, Y, Sun, F, Wang, L, Gao, M, Xie, Y, Sun, Y, Liu, H, Yuan, Y, Yi, W, and Huang, Z (2020): Virus-induced p38 MAPK activation facilitates viral infection. Theranostics 10, 12223.

Choi, M-S, Heo, J, Yi, C-M, Ban, J, Lee, N-J, Lee, N-R, Kim, SW, Kim, N-J, and Inn, K-S (2016): A novel p38 mitogen activated protein kinase (MAPK) specific inhibitor suppresses respiratory syncytial virus and influenza A virus replication by inhibiting virus-induced p38 MAPK activation. Biochemical and biophysical research communications 477, 311–316.

Cohen, P (1997): The search for physiological substrates of MAP and SAP kinases in mammalian cells. Trends in cell biology 7, 353–361.

Cook, M (2016): The role of MAPK p38 stress pathway-induced cellular translation in human and macaque cells targeted during B virus infection.

Cuadrado, A, and Nebreda, AR (2010): Mechanisms and functions of p38 MAPK signalling. Biochem J 429, 403–17.

Du, Q, Huang, Y, Wang, T, Zhang, X, Chen, Y, Cui, B, Li, D, Zhao, X, Zhang, W, Chang, L, and Tong, D (2016): Porcine circovirus type 2 activates PI3K/Akt and p38 MAPK pathways to promote interleukin-10 production in macrophages via Cap interaction of gC1qR. Oncotarget 7, 17492–507.

Food, U, and Administration, D (2020): Coronavirus (COVID-19) update: FDA issues emergency use authorization for potential COVID-19 treatment. FDA news release.

Galván Morales, MÁ, Cabello Gutiérrez, C, Mejía Nepomuceno, F, Valle Peralta, L, Valencia Maqueda, E, and Manjarrez Zavala, ME (2014): Parainfluenza virus type 1 induces epithelial IL-8 production via p38-MAPK signalling. Journal of immunology research 2014.

Griego, SD, Weston, CB, Adams, JL, Tal-Singer, R, and Dillon, SB (2000): Role of p38 mitogen-activated protein kinase in rhinovirus-induced cytokine production by bronchial epithelial cells. The Journal of Immunology 165, 5211–5220.

Henklova, P, Vrzal, R, Papouskova, B, Bednar, P, Jancova, P, Anzenbacherova, E, Ulrichova, J, Maurel, P, Pavek, P, and Dvorak, Z (2008): SB203580, a pharmacological inhibitor of p38 MAP kinase transduction pathway activates ERK and JNK MAP kinases in primary cultures of human hepatocytes. European journal of pharmacology 593, 16–23.

Higgins, CA, Nilsson-Payant, BE, Kurland, A, Adhikary, P, Golynker, I, Danziger, O, Panis, M, Rosenberg, BR, and Johnson, JR (2021): SARS-CoV-2 hijacks p38β/MAPK11 to promote viral protein translation. bioRxiv.

Hopcraft, SE, and Evans, MJ (2015): Selection of a hepatitis C virus with altered entry factor requirements reveals a genetic interaction between the E1 glycoprotein and claudins. Hepatology 62, 1059–69.

Hovi, T, Järvinen, A, Pyhälä, R, Ristola, M, and Salminen, M (2002): Viruses and antiviral drug resistance. Duodecim; laaketieteellinen aikakauskirja 118, 911–918.

Huang, G, Shi, LZ, and Chi, H (2009): Regulation of JNK and p38 MAPK in the immune system: signal integration, propagation and termination. Cytokine 48, 161–169.

Irwin, KK, Renzette, N, Kowalik, TF, and Jensen, JD (2016): Antiviral drug resistance as an adaptive process. Virus evolution 2, vew014.

Khandelwal, N, Chander, Y, Kumar, R, Nagori, H, Verma, A, Mittal, P, Kamboj, S, Verma, SS, Khatreja, S, and Pal, Y (2021): Studies on growth characteristics and cross-neutralization of wild-type and delta SARS-CoV-2 from Hisar (India). Frontiers in Cellular and Infection Microbiology 11, 771524.

Koul, HK, Pal, M, and Koul, S (2013): Role of p38 MAP kinase signal transduction in solid tumors. Genes & cancer 4, 342–359.

Kumar, N, Barua, S, Riyesh, T, Chaubey, KK, Rawat, KD, Khandelwal, N, Mishra, AK, Sharma, N, Chandel, SS, and Sharma, S (2016): Complexities in isolation and purification of multiple viruses from mixed viral infections: viral interference, persistence and exclusion. PloS one 11, e0156110.

Kumar, N, Khandelwal, N, Kumar, R, Chander, Y, Rawat, KD, Chaubey, KK, Sharma, S, Singh, SV, Riyesh, T, and Tripathi, BN (2019a): Inhibitor of sarco/endoplasmic reticulum calcium-ATPase impairs multiple steps of paramyxovirus replication. Frontiers in Microbiology 10, 209.

Kumar, N, Khandelwal, N, Kumar, R, Chander, Y, Rawat, KD, Chaubey, KK, Sharma, S, Singh, SV, Riyesh, T, Tripathi, BN, and Barua, S (2019b): Inhibitor of Sarco/Endoplasmic Reticulum Calcium-ATPase Impairs Multiple Steps of Paramyxovirus Replication. Front Microbiol 10, 209.

Kumar, N, Liang, Y, Parslow, TG, and Liang, Y (2011a): Receptor tyrosine kinase inhibitors block multiple steps of influenza a virus replication. J Virol 85, 2818–27.

Kumar, N, Maherchandani, S, Kashyap, SK, Singh, SV, Sharma, S, Chaubey, KK, and Ly, H (2014): Peste des petits ruminants virus infection of small ruminants: a comprehensive review. Viruses 6, 2287–327.

Kumar, N, Sharma, NR, Ly, H, Parslow, TG, and Liang, Y (2011b): Receptor tyrosine kinase inhibitors that block replication of influenza a and other viruses. Antimicrob Agents Chemother 55, 5553–9.

Kumar, N, Sharma, S, Kumar, R, Tripathi, BN, Barua, S, Ly, H, and Rouse, BT (2020): Host-Directed Antiviral Therapy. Clin Microbiol Rev 33.

Kumar, N, Xin, ZT, Liang, Y, Ly, H, and Liang, Y (2008): NF-kappaB signaling differentially regulates influenza virus RNA synthesis. J Virol 82, 9880–9.

Kumar, R, Afsar, M, Khandelwal, N, Chander, Y, Riyesh, T, Dedar, RK, Gulati, BR, Pal, Y, Barua, S, and Tripathi, BN (2021): Emetine suppresses SARS-CoV-2 replication by inhibiting interaction of viral mRNA with eIF4E. Antiviral research 189, 105056.

Kumar, R, Khandelwal, N, Chander, Y, Nagori, H, Verma, A, Barua, A, Godara, B, Pal, Y, Gulati, BR, and Tripathi, BN (2022): S-adenosylmethionine-dependent methyltransferase inhibitor DZNep blocks transcription and translation of SARS-CoV-2 genome with a low tendency to select for drug-resistant viral variants. Antiviral research 197, 105232.

Kumar, R, Khandelwal, N, Chander, Y, Riyesh, T, Tripathi, BN, Kashyap, SK, Barua, S, Maherchandani, S, and Kumar, N (2018a): MNK1 inhibitor as an antiviral agent suppresses buffalopox virus protein synthesis. Antiviral research 160, 126–136.

Kumar, R, Khandelwal, N, Chander, Y, Riyesh, T, Tripathi, BN, Kashyap, SK, Barua, S, Maherchandani, S, and Kumar, N (2018b): MNK1 inhibitor as an antiviral agent suppresses buffalopox virus protein synthesis. Antiviral Res 160, 126–136.

Kumar, R, Khandelwal, N, Thachamvally, R, Tripathi, BN, Barua, S, Kashyap, SK, Maherchandani, S, and Kumar, N (2018c): Role of MAPK/MNK1 signaling in virus replication. Virus research 253, 48–61.

Kumar, R, Khandelwal, N, Thachamvally, R, Tripathi, BN, Barua, S, Kashyap, SK, Maherchandani, S, and Kumar, N (2018d): Role of MAPK/MNK1 signaling in virus replication. Virus Res 253, 48–61.

Lamarche, MJ, Borawski, J, Bose, A, Capacci-Daniel, C, Colvin, R, Dennehy, M, Ding, J, Dobler, M, Drumm, J, Gaither, LA, Gao, J, Jiang, X, Lin, K, McKeever, U, Puyang, X, Raman, P, Thohan, S, Tommasi, R, Wagner, K, Xiong, X, Zabawa, T, Zhu, S, and Wiedmann, B (2012): Anti-hepatitis C virus activity and toxicity of type III phosphatidylinositol-4-kinase beta inhibitors. Antimicrob Agents Chemother 56, 5149–56.

Lee, N, Wong, CK, Chan, PK, Lun, SW, Lui, G, Wong, B, Hui, DS, Lam, CW, Cockram, CS, Choi, KW, Yeung, AC, Tang, JW, and Sung, JJ (2007): Hypercytokinemia and hyperactivation of phospho-p38 mitogen-activated protein kinase in severe human influenza A virus infection. Clin Infect Dis 45, 723–31.

Li, DK, and Chung, RT (2019): Overview of direct-acting antiviral drugs and drug resistance of hepatitis C virus. Hepatitis C Virus Protocols, 3–32.

Locarnini, S, and Bowden, S (2010): Drug resistance in antiviral therapy. Clinics in liver disease 14, 439–459.

Maik-Rachline, G, Lifshits, L, and Seger, R (2020): Nuclear P38: roles in physiological and pathological processes and regulation of nuclear translocation. International journal of molecular sciences 21, 6102.

Merino-Ramos, T, Vázquez-Calvo, Á, Casas, J, Sobrino, F, Saiz, J-C, and Martín-Acebes, MA (2016): Modification of the host cell lipid metabolism induced by hypolipidemic drugs targeting the acetyl coenzyme A carboxylase impairs West Nile virus replication. Antimicrobial agents and chemotherapy 60, 307–315.

Mikkelsen, SS, Jensen, SB, Chiliveru, S, Melchjorsen, J, Julkunen, I, Gaestel, M, Arthur, JS, Flavell, RA, Ghosh, S, and Paludan, SR (2009): RIG-I-mediated activation of p38 MAPK is essential for viral induction of interferon and activation of dendritic cells: dependence on TRAF2 and TAK1. J Biol Chem 284, 10774–82.

Mudaliar, P, Pradeep, P, Abraham, R, and Sreekumar, E (2021): Targeting cap-dependent translation to inhibit Chikungunya virus replication: selectivity of p38 MAPK inhibitors to virus-infected cells due to autophagy-mediated down regulation of phospho-ERK. Journal of General Virology 102, 001629.

Pawlotsky, JM (2012): The science of direct-acting antiviral and host-targeted agent therapy. Antivir Ther 17, 1109–17.

Peng, H, Shi, M, Zhang, L, Li, Y, Sun, J, Zhang, L, Wang, X, Xu, X, Zhang, X, and Mao, Y (2014): Activation of JNK1/2 and p38 MAPK signaling pathways promotes enterovirus 71 infection in immature dendritic cells. BMC microbiology 14, 1–9.

Pillay, D, and Zambon, M (1998): Antiviral drug resistance. Bmj 317, 660–662.

Rajnik, M, Cascella, M, Cuomo, A, Dulebohn, SC, and Di Napoli, R (2021): Features, evaluation, and treatment of coronavirus (COVID-19). Uniformed Services University Of The Health Sciences.

Shapiro, L, Heidenreich, KA, Meintzer, MK, and Dinarello, CA (1998): Role of p38 mitogen-activated protein kinase in HIV type 1 production in vitro. Proceedings of the National Academy of Sciences 95, 7422–7426.

Shepherd, FA, Rodrigues Pereira, J, Ciuleanu, T, Tan, EH, Hirsh, V, Thongprasert, S, Campos, D, Maoleekoonpiroj, S, Smylie, M, Martins, R, van Kooten, M, Dediu, M, Findlay, B, Tu, D, Johnston, D, Bezjak, A, Clark, G, Santabarbara, P, Seymour, L, and National Cancer Institute of Canada Clinical Trials, G (2005): Erlotinib in previously treated non-small-cell lung cancer. N Engl J Med 353, 123–32.

Su, A-r, Qiu, M, Li, Y-l, Xu, W-t, Song, S-w, Wang, X-h, Song, H-y, Zheng, N, and Wu, Z-w (2017): BX-795 inhibits HSV-1 and HSV-2 replication by blocking the JNK/p38 pathways without interfering with PDK1 activity in host cells. Acta Pharmacologica Sinica 38, 402–414.

Sugasti-Salazar, M, Llamas-González, YY, Campos, D, and González-Santamaría, J (2021): Inhibition of p38 Mitogen-Activated Protein Kinase Impairs Mayaro Virus Replication in Human Dermal Fibroblasts and HeLa Cells. Viruses 13, 1156.

Team, C-I, Kujawski, SA, Wong, KK, Collins, JP, Epstein, L, Killerby, ME, Midgley, CM, Abedi, GR, Ahmed, NS, and Almendares, O (2020): First 12 patients with coronavirus disease 2019 (COVID-19) in the United States. MedRxiv, 2020.03. 09.20032896.

ul Qamar, MT, Alqahtani, SM, Alamri, MA, and2 Chen, L-L (2020): Structural basis of SARS-CoV-2 3CLpro and anti-COVID-19 drug discovery from medicinal plants. Journal of pharmaceutical analysis 10, 313–319.

Venter, JC, Adams, MD, Myers, EW, Li, PW, Mural, RJ, Sutton, GG, Smith, HO, Yandell, M, Evans, CA, and Holt, RA (2001): The sequence of the human genome. Science 291, 1304–1351.

Warren, TK, Jordan, R, Lo, MK, Ray, AS, Mackman, RL, Soloveva, V, Siegel, D, Perron, M, Bannister, R, and Hui, HC (2016): Therapeutic efficacy of the small molecule GS-5734 against Ebola virus in rhesus monkeys. Nature 531, 381–385.

Watanabe, K, Takizawa, N, Katoh, M, Hoshida, K, Kobayashi, N, and Nagata, K (2001): Inhibition of nuclear export of ribonucleoprotein complexes of influenza virus by leptomycin B. Virus research 77, 31–42.

Xu, X, Miao, J, Shao, Q, Gao, Y, and Hong, L (2020): Apigenin suppresses influenza A virus□induced RIG□I activation and viral replication. Journal of Medical Virology 92, 3057–3066.

Zeisel, MB, Lupberger, J, Fofana, I, and Baumert, TF (2013): Host-targeting agents for prevention and treatment of chronic hepatitis C - perspectives and challenges. J Hepatol 58, 375–84.

Zhan, Y, Yu, S, Yang, S, Qiu, X, Meng, C, Tan, L, Song, C, Liao, Y, Liu, W, and Sun, Y (2020): Newcastle Disease virus infection activates PI3K/Akt/mTOR and p38 MAPK/Mnk1 pathways to benefit viral mRNA translation via interaction of the viral NP protein and host eIF4E. PLoS pathogens 16, e1008610.

Zhao, Z, Li, H, Wu, X, Zhong, Y, Zhang, K, Zhang, Y-P, Boerwinkle, E, and Fu, Y-X (2004): Moderate mutation rate in the SARS coronavirus genome and its implications. BMC evolutionary biology 4, 1–9

